# HiCrayon reveals distinct layers of multi-state 3D chromatin organization

**DOI:** 10.1101/2024.02.11.579821

**Authors:** Ben Nolan, Hannah L. Harris, Achyuth Kalluchi, Timothy E. Reznicek, Christopher T. Cummings, M. Jordan Rowley

## Abstract

The co-visualization of chromatin conformation with 1D ‘omics data is key to the multi-omics driven data analysis of 3D genome organization. Chromatin contact maps are often shown as 2D heatmaps and visually compared to 1D genomic data by simple juxtaposition. While common, this strategy is imprecise, placing the onus on the reader to align features with each other. To remedy this, we developed HiCrayon, an interactive tool that facilitates the integration of 3D chromatin organization maps and 1D datasets. This visualization method integrates data from genomic assays directly into the chromatin contact map by coloring interactions according to 1D signal. HiCrayon is implemented using R shiny and python to create a graphical user interface (GUI) application, available in both web or containerized format to promote accessibility. HiCrayon is implemented in R, and includes a graphical user interface (GUI), as well as a slimmed-down web-based version that lets users quickly produce publication-ready images. We demonstrate the utility of HiCrayon in visualizing the effectiveness of compartment calling and the relationship between ChIP-seq and various features of chromatin organization. We also demonstrate the improved visualization of other 3D genomic phenomena, such as differences between loops associated with CTCF/cohesin vs. those associated with H3K27ac. We then demonstrate HiCrayon’s visualization of organizational changes that occur during differentiation and use HiCrayon to detect compartment patterns that cannot be assigned to either A or B compartments, revealing a distinct 3rd chromatin compartment. Overall, we demonstrate the utility of co-visualizing 2D chromatin conformation with 1D genomic signals within the same matrix to reveal fundamental aspects of genome organization.

Local version: https://github.com/JRowleyLab/HiCrayon

Web version: https://jrowleylab.com/HiCrayon

## 1 Background

The study of 3D genome organization relies on integrating data from multiple sources, including the co-analysis of 2D maps of chromatin conformation with 1D genomic signals such as ChIP-seq tracks for histone modifications or architectural proteins. Genome-wide chromatin conformation capture experiments, such as Hi-C, Micro-C, and Pore-C, measure long-range chromatin interactions, which are most commonly represented as 2D heatmaps [1–3]. These chromatin contact maps detail several features that have revealed fundamental principles of chromatin organization [4]. Some of the earliest maps describe the propensity of loci to interact in A (active) and B (inactive) compartments [1], with more recent work showing that this segregation can occur at kilobase scale [5, 6]. Genome-wide maps of chromatin organization also reveal high-intensity punctate signals corresponding to CTCF loops in mammalian cells [7] and Polycomb (Pc) loops in *Drosophila melanogaster* [8–11]. Many also find it useful to examine domains of interactions, sometimes referred to as Topologically Associated Domains (TADs), which appear as triangles of intense signal near the diagonal [12–15]. We should note, however, that interaction domains can sometimes describe different features depending on the context (e.g. CTCF loop domains vs. compartment domains) [4, 7, 16–18].

Our understanding of 3D genome organization is often derived from the comparison of 2D chromatin contact maps with 1D genomic datasets. For example, broad-scale comparisons of the plaid-like Hi-C compartment pattern with histone modification ChIP-seq tracks reveal the propensity for chromatin with active histone modifications to locate in the A compartment [1]. When viewed at fine scale, we recently showed that localization to the A compartment is a precise, fundamental characteristic of active enhancers and promoters [5]. Another well-known example is the overlap between 2D punctate loops and 1D CTCF ChIP-seq data, revealing the overwhelming presence of CTCF at these loop anchors [7]. In contrast, a comparison of high-intensity 2D punctate loops with 1D ChIP-seq signal revealed a lack of CTCF occupancy in *D. melanogaster* [16]. Instead, *D. melanogaster* loop anchors are occupied by Pc and Pipsqeuak [8–11].

Currently, the visual comparison of 2D and 1D genomic datasets at example loci is most often accomplished through the juxtaposition of heatmap and signal tracks (Figure 1A) [19, 20]. While useful, this simple visualization relies on “eyeballing” the relationships, with the user visually aligning peaks of 1D signal with 2D features. Naturally, this current practice can be imprecise and prone to human visual biases. To aid the precise visualization of overlapping 2D and 1D features, we developed HiCrayon for the visual integration of 1D genomic signals with 2D chromatin contacts within a single matrix. We showcase the capabilities of HiCrayon by coloring chromatin contact maps by ChIP-seq signal to reveal features, including CTCF and non-CTCF associated loops, as well as the changes to chromatin organization that occur during differentiation. We also demonstrate the ability of HiCrayon to reveal distinct segregation of chromatin into multiple distinct compartment states, which the eigenvector and its commonly accepted binary ‘A’ and ‘B’ designation fails to describe. We then demonstrate the ability of HiCrayon to depict distinct interactions associated with CTCF loops and fine-scale compartments, allowing one to distinguish interaction domains associated with each. These results demonstrate the ability of HiCrayon to perform advanced data visualization to uncover fundamental principles of 3D chromatin organization.

**Fig. 1.**
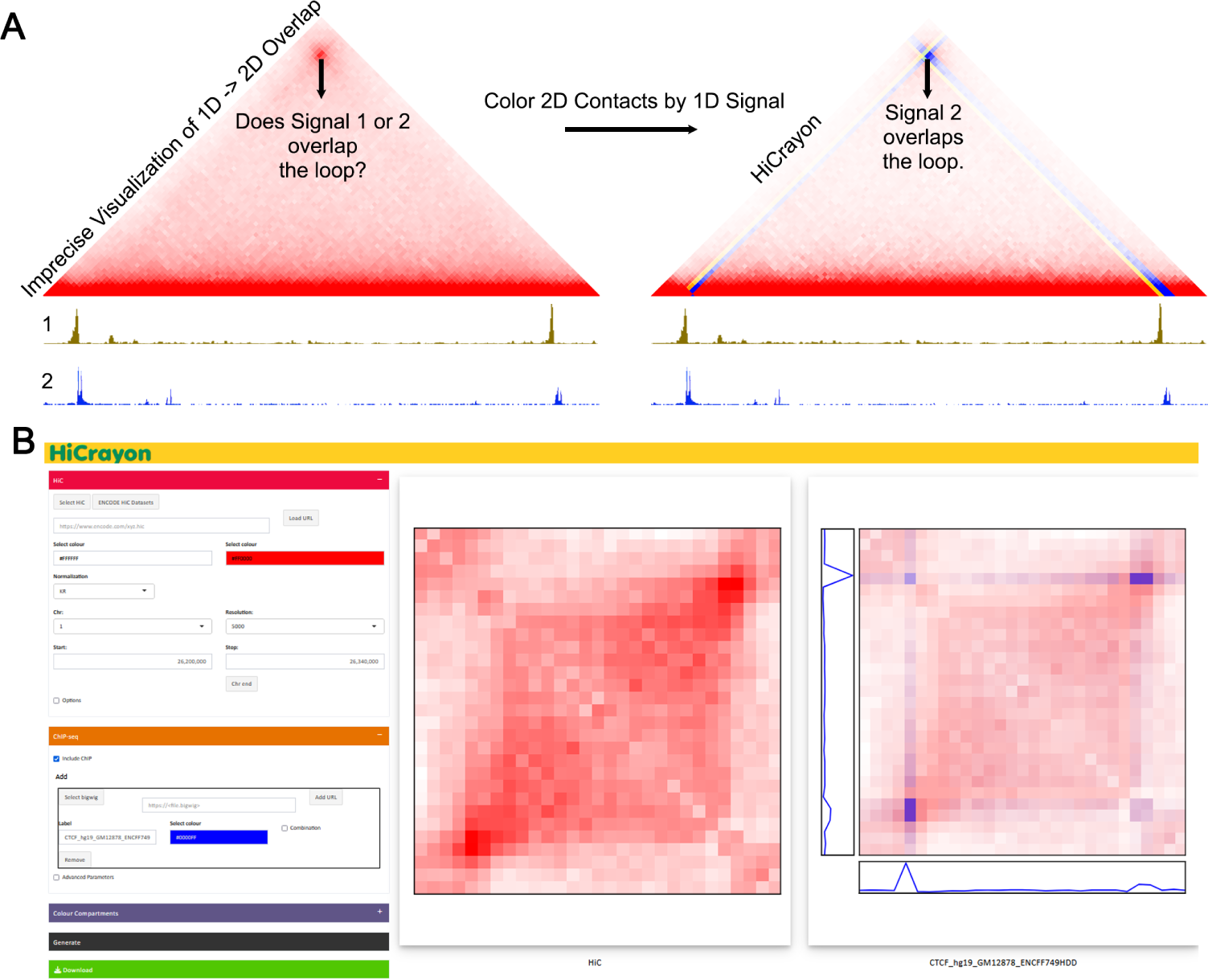
HiCrayon. **A:** A common visualization method juxtaposes 1-dimensional (1D) tracks under a 2-dimensional (2D) map of chromatin contacts (left). However, it can be difficult to determine which 1D feature overlaps the 2D feature of interest. HiCrayon creates a 2D contact matrix where the 2D contacts are color-coded by 1D signal (right). B: A screenshot of HiCrayon’s graphical user interface (GUI).

## 2 Results

### 2.1 HiCrayon Overview

To overcome limitations in the comparison of 1D and 2D chromatin data, we developed HiCrayon for coloring chromatin contact maps by 1D signal (Figure 1A). HiCrayon is implemented using a combination of Python and the R shiny package [21] for a menu-driven graphical user interface (GUI) (Figure 1B). The application is available in two forms, 1) A full-functionality, local version installed through Github, https://github.com/JRowleyLab/HiCrayon, that allows the visualization of files stored locally or via a URL; or 2) a website version, https://jrowleylab.com/HiCrayon, with more limited functionality to allow ease of access for users who wish to visualize previously published datasets via URL, such as those stored on the ENCODE data portal [22, 23]. Images generated by HiCrayon can be downloaded in SVG and PNG format.

HiCrayon allows the selection of multiple 1D bigwig signal tracks to be overlayed at the selected 2D locus. Visualization is accomplished by 1D to 2D transformation of signals, weighted by the interactions matrix along with linear interpolation of color matrices (see Methods). Advanced options allow users to adjust the degree of overlay between the 1D-to-2D signal matrix on the 2D chromatin contact matrix. Each 1D- to-2D signal matrix can be visualized separately or can be overlaid with each other within a single, fully customizable bespoke color-blended image. HiCrayon allows visualization of 1D signal with negative and positive values, as commonly obtained from compartment identification [24]. In this mode, HiCrayon uses two colors to distinguish positive from negative values, thereby allowing distinct visualization of A and B compartment interactions.

### 2.2 HiCrayon improves the visualization of protein occupancy overlap at punctate loops

Maps of 3D chromatin organization in mammals have revealed thousands of punctate chromatin loops [7, 25, 26]. The most prominent loops are formed by cohesin-mediated extrusion, which is blocked at CTCF loop anchors [4, 27–32]. Therefore, punctate loops in mammals are typically enriched for CTCF and RAD21, a member of the cohesin complex [7, 33–36]. We also recently showed that CTCF loops are often comprised of fairly diffuse interactions when viewed at high resolution [5] (Figure 2A, left panel). Using this high-resolution Hi-C map in lymphoblastoid cell lines (LCLs) [5], we colored contacts by HiCrayon to visualize the overlap between chromatin interactions and published ChIP-seq for CTCF (red) and RAD21 (blue). We found that HiCrayon at high resolution can help visualize the center of loop foci where an intense Hi-C signal coincides with CTCF and cohesin together (purple) (Figure 2A). We find that this strategy aids with the visualization of overlapping features such as CTCF and cohesin at loop anchors (Figure 2A) as well as distinguishes non-overlapping features, such as specific H3K27ac (yellow) sites that, at this locus, do not coincide with CTCF loop anchors (Figure 2B). Because H3K27ac does not correspond to a strong interaction signal within this region, this example also demonstrates that HiCrayon avoids coloring non-interacting features even when a 1D signal is present (Figure 2B). In contrast, by simultaneously coloring the Hi-C map with three colors, we found that HiCrayon was able to visually distinguish H3K27ac interactions from that of CTCF/cohesin loops (Figure 2C,D).

**Fig. 2.**
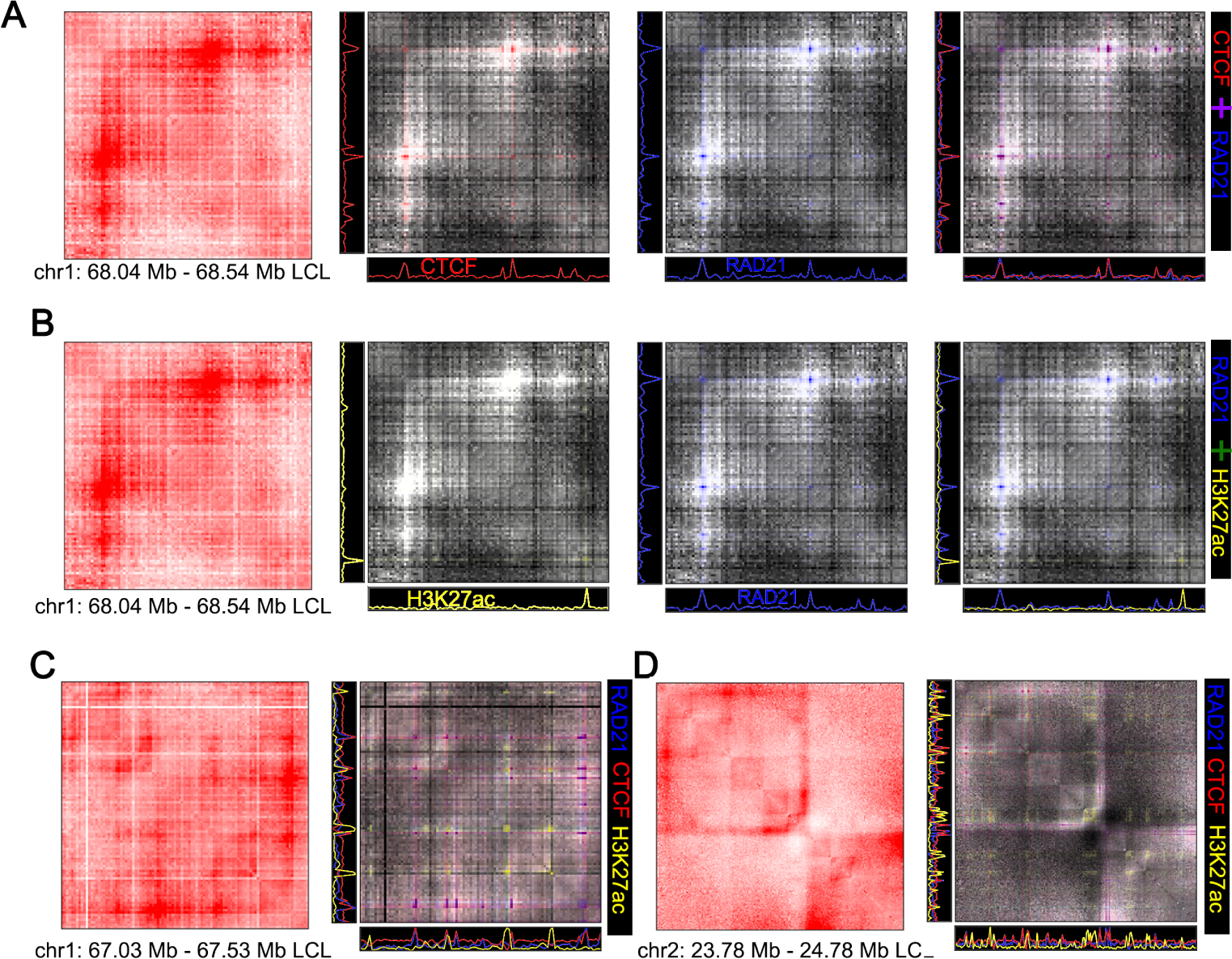
H3K27ac forms loops distinct from that of CTCF and cohesin. **A:** Example of a loop occupied by both CTCF and cohesin. Left to right: Distance normalized Hi-C in LCLs, colored by ChIP-seq signal for CTCF (red), cohesin (blue), or both. Linear interpolation of colors makes the overlap between CTCF and cohesin at loop anchors visible as purple. **B:** The same loop as in A, but using H3K27ac ChIP-seq signal to show how non-overlapping features are not colored. **C:** and **D:** 3-color HiCrayon images for two different loci showing H3K27ac-specific interactions (yellow) that are distinct from CTCF / cohesin loops.

We then used HiCrayon to visualize punctate loops that have been found in Hi-C maps of *D. melanogaster* Kc167 cells [8, 10]. In contrast to the thousands of punctate CTCF loops found in mammals, there are only a few hundred intense punctate loops in Hi-C maps of interphase *D. melanogaster* [8–10, 16]. While these punctate signals are commonly referred to as Polycomb (Pc) loops [9, 10], the anchors are also occupied by the architectural protein Pipsqueak (Psq) [11] (Figure 3A). Zooming in on the loop at 250 bp resolution, HiCrayon captures the overlap (pink) between Pc (purple) and Psq (yellow) (Figure 3B). In contrast, HiCrayon coloring found that, despite high H3K27me3 levels in the vicinity, these interactions precisely overlap a peak of H3K27ac (yellow) and not H3K27me3 (purple) (Figure 3C) [11]. This indicates that, despite the overlap with Pc, these Psq-associated loops correspond to islands of active H3K27acassociated chromatin in the midst of the broader H3K27me3 repressive chromatin [11]. Overall, we find that HiCrayon has the ability to visually identify precise overlaps between protein occupancy and features of 3D chromatin organization.

**Fig. 3.**
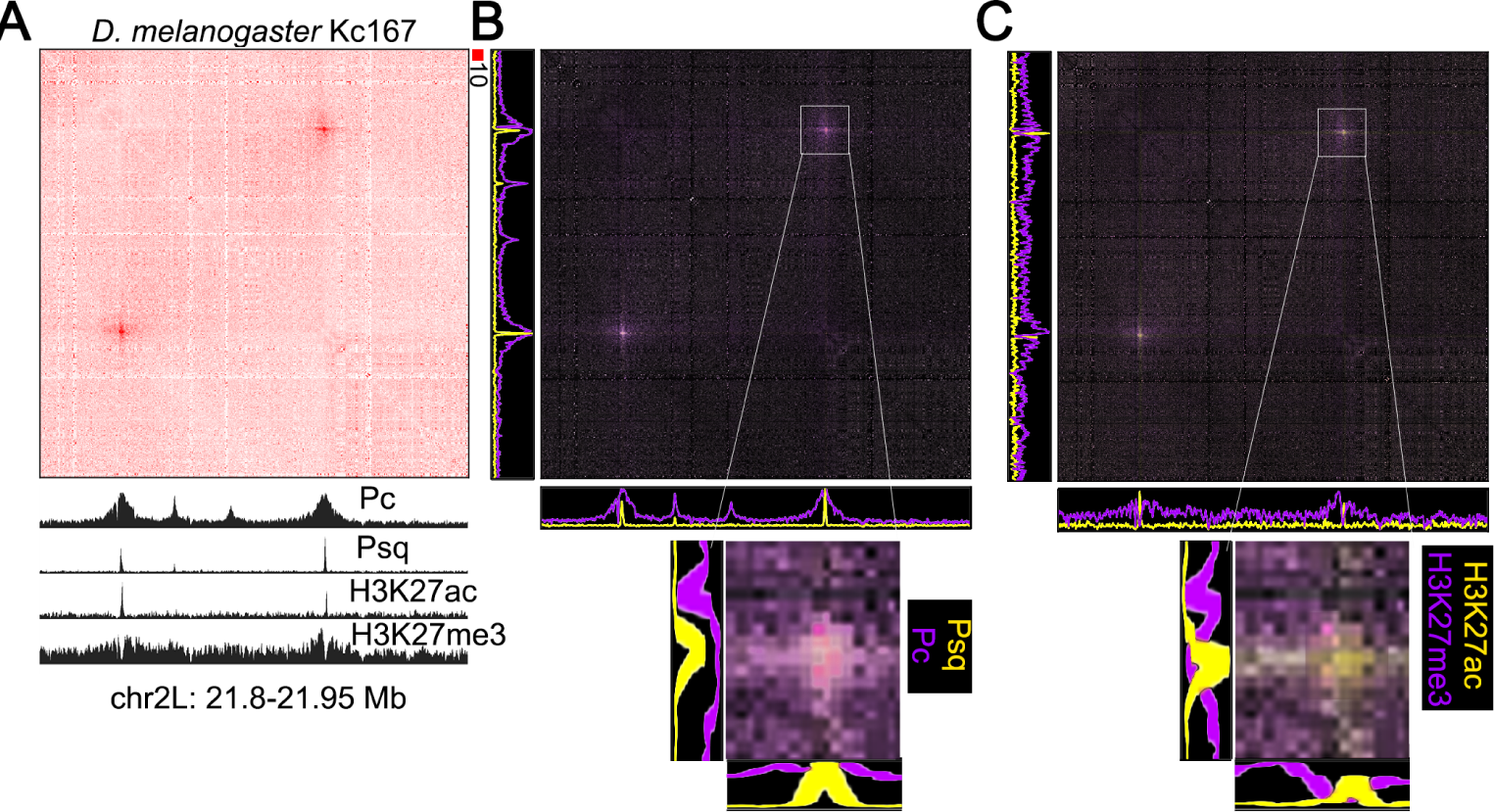
Punctate loops in *Drosophila* are associated with Pc, Psq, and H3K27ac. **A-C:** Example of a punctate loop found in *Drosophila melanogaster* Kc167 cells shown by A) distance normalized Hi-C with 1D tracks representing ChIP-seq signal for Polycomb (Pc), Pipsqueak (Psq), H3K27ac, and H3K27me3; B) HiCrayon coloring of interactions by Psq (yellow) and Pc (purple) ChIP-seq signal, resulting in pink at overlapping features; C) HiCrayon coloring by H3K27ac (yellow) and H3K27me3 (purple) ChIP-seq signal. The yellow signal at the loop reflects the peak in H3K27ac and dip in H3K27me3 signal precisely at the loop anchors.

### 2.3 HiCrayon visualizes overlapping changes to protein occupancy and genome organization during differentiation

Several studies have profiled changes in genome organization that occur during differentiation [37–40]. We asked if HiCrayon is able to visualize differential loops corresponding to altered occupancy at anchors in mouse embryonic stem cells (mESC), and neural progenitor cells (NPC) [39]. Indeed, HiCrayon maps of CTCF (pink) and H3K27ac (yellow) highlighted the altered structure of an example locus (Figure 4A). Specifically, HiCrayon reveals a loss of a short loop near the pluripotency marker *Sox2*, along with a gain in a longer loop after differentiation to NPCs (Figure 4A). In each case, we found that H3K27ac was nearby but not directly at the CTCF loop anchors (Figure 4A), an important distinction that would be missed by simple juxtaposition.

**Fig. 4.**
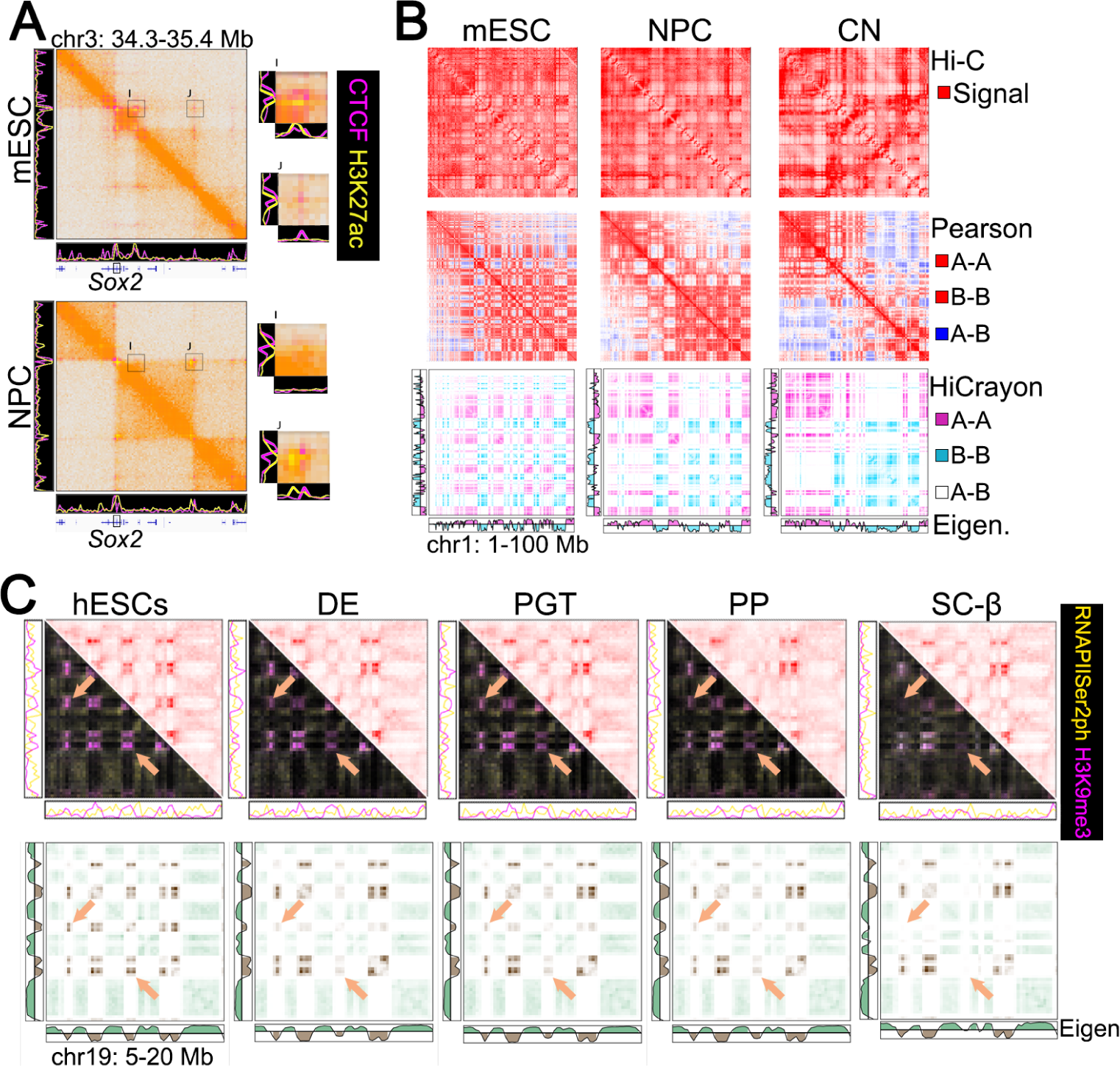
HiCrayon’s compartment visualization reveals coordinated changes in histone marks and compartments during differentiation. **A:** Example locus with a differential loop that is altered during differentiation from mouse embryonic stem cells (ES) and neural progenitor cells (NPC). HiCrayon coloring by CTCF (pink) and H3K27ac (yellow) helps illustrate the changes that occur primarily across two loci, I and J, which correspond to two CTCF loop anchors. **B:** Comparison of Hi-C maps, Pearson correlation matrices, and HiCrayon to color compartments and changes that occur during differentiation of mouse embryonic stem cells (mESCs), neural progenitor cells (NPCs), and cortical neurons (CN). The color legend on the right indicates how features are colored in each visualization method. **C:** Top: Hi-C and HiCrayon maps colored by RNA Polymerase Serine 2 phosphorylation (RNAPIISer2ph) (yellow) and H3K9me3 (purple) across differentiating cells. Bottom: The same contact maps, but using HiCrayon to color by the compartment eigenvector. Arrows highlight regions with decreased B-B interactions.

Next, we tested if HiCrayon could be used to visualize changes in compartments that occur during differentiation into cortical neurons (CN) [39]. A and B compartments are represented by a plaid-like pattern in Hi-C maps (Figure 4B, top), and the Pearson correlation matrix is often used to visualize this pattern (Figure 4B, middle). While this strategy helps distinguish A-B interactions (blue) from A-A or B-B (red), it fails to distinguish between A-A and B-B interactions themselves, as they are both represented by the same color (Figure 4B, middle). To facilitate the distinct visualization of A-A and B-B interactions, we built into HiCrayon the ability to color the Hi-C map by the eigenvector. Coloring interactions by the eigenvector provides a visual distinction of A and B compartment interactions and highlights differences between maps (Figure 4B, bottom). Indeed, we found that HiCrayon in mESC, NPC, and CN highlights the dramatic reorganization of compartments that occurs during differentiation (Figure 4B).

Recently, Hi-C maps during pancreatic islet differentiation revealed changes to chromatin organization [38]. Differentiation of human embryonic stem cells (hESC) to that of definitive endoderm (DE), primitive gut tube (PGT), pancreatic progenitor (PP), and stem cell-derived *β*-cells (sc-*β*) results in altered compartments associated with a loss of H3K9me3 at some loci [38]. We used HiCrayon to color by RNAPIISer2ph (yellow) and H3K9me3 (purple), which allowed direct visual identification of these changes, which corresponded to changes in H3K9me3 (Figure 4C, top). Further visualization, by coloring these maps by the eigenvector, allowed direct representation of the decreased B-B interactions precisely at loci that lose H3K9me3 during pancreatic differentiation (Figure 4C, bottom).

### 2.4 HiCrayon visually distinguishes a 3rd chromatin compartment

Annotation of A/B compartments has proven informative for numerous studies of 3D genome organization (reviewed in [24]). However, recent work suggests that this binary segregation represents an oversimplification and that compartments are rather more complicated than two states and may even be organized into three or more distinct compartments or subcompartments [7, 24, 41, 42]. Most representations of chromatin contact maps use a single gradient color scale (e.g., white to red), making it difficult to visually detect multiple states (Figure 5A). Using HiCrayon to color interactions by the A/B compartment eigenvector, we noticed that some interactions failed to be annotated as A or B, i.e., where the eigenvector approaches zero (Figure 5B). We found that these loci have compartment-like patterns distinct from the regions that the eigenvector identified as A or B (Figure 5C). To determine the chromatin status of these loci, we used HiCrayon to color the Hi-C map by histone modification ChIP-seq signal. While H3K27ac (green) overlaps the A-A compartment pattern, and H3K9me3 (purple) overlaps the B-B compartment pattern on this chromosome, we noticed that H3K27me3 (orange) signal is distinct from the others (Figure 5D and E). Furthermore, we observe that bins are remarkably dominant in a single histone mark, in comparison to the other two histone marks (Figure 5F). These results provide evidence for a more complex model of compartment organization than what is depicted by eigenvector-based compartment annotation.

**Fig. 5.**
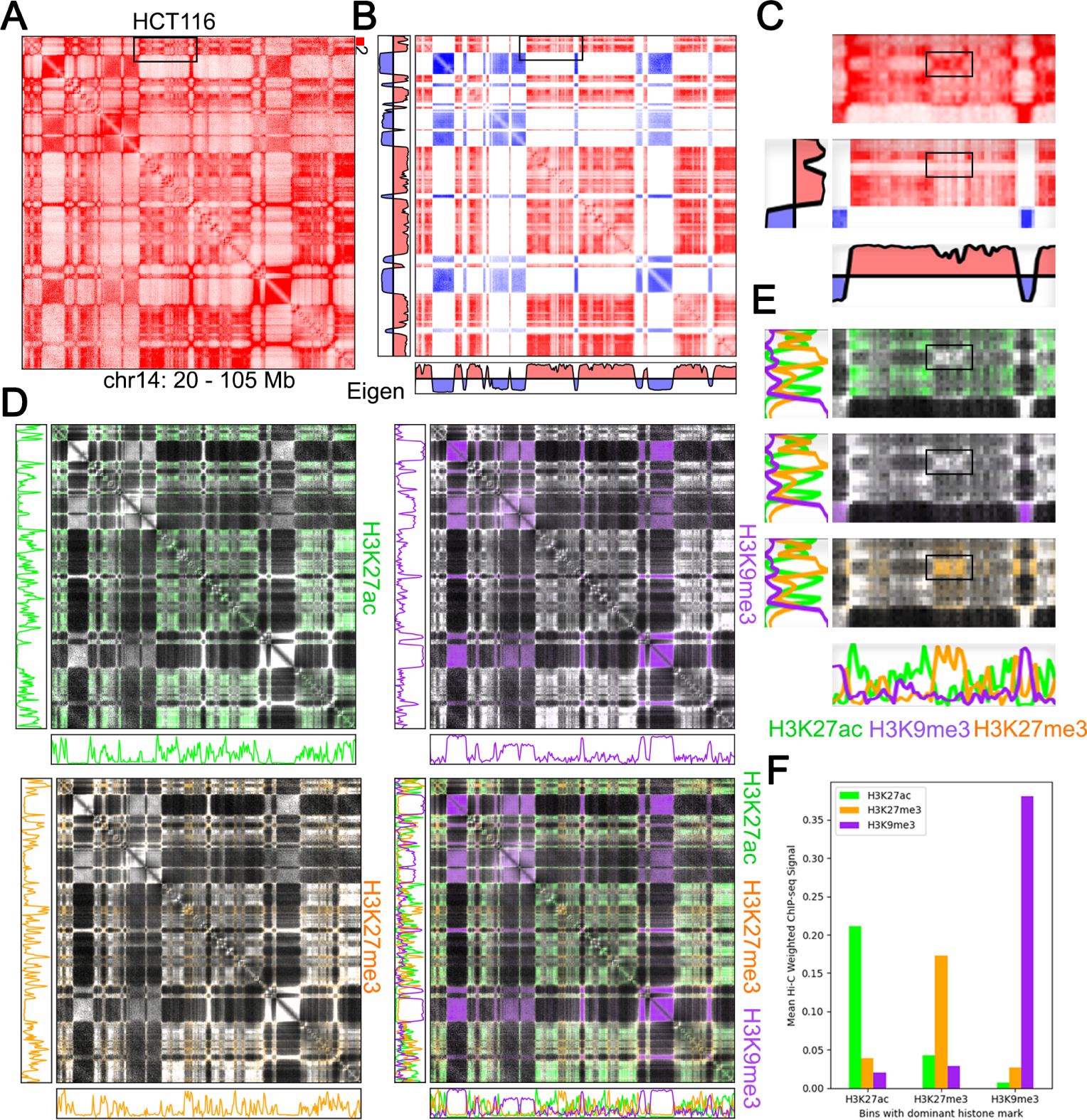
HiCrayon visualization reveals more than two compartments. **A-C:** Hi-C map (A), HiCrayon coloring by the eigenvector (B), and zoomed-in region (C) depicting a strong compartmental pattern on chr14 of HCT116 cells, that cannot be assigned as either A or B. **D,E:** HiCrayon coloring of the compartment interactions by H3K27ac (green), H3K9me3 (purple), or H3K27me3 (orange). **F:** Comparison of mean Hi-C weighted ChIP-seq signal for H3K27ac, H3K27me3, and H3K9me3 in each bin showcasing a strong mutual exclusivity of histone modifications.

### 2.5 Chromatin interaction domains are formed by distinct layers of genome organization

When we colored Hi-C maps by the A/B compartment eigenvector in *D. melanogaster* Kc167 cells, we noticed that it colored domain-like squares around the diagonal, often referred to as Topologically Associated Domains (TADs) (Figure 6A). As previously reported, *D. melanogaster* lacks CTCF loops [16], and as such, we’ve found that the TADs correspond exceptionally well with compartment intervals, with A and B compartments represented by red and blue respectively (Figure 6A) [16], albeit other factors likely also influence domain structures [16, 43–45]. Indeed, HiCrayon visualization of H3K27me3 and H3K27ac exposes the alternating chromatin activity states within the region and demonstrates the close relationship between chromatin state, compartment intervals, and domains in *D. melanogaster* (Figure 6A) [13, 14, 16, 44, 46].

**Fig. 6.**
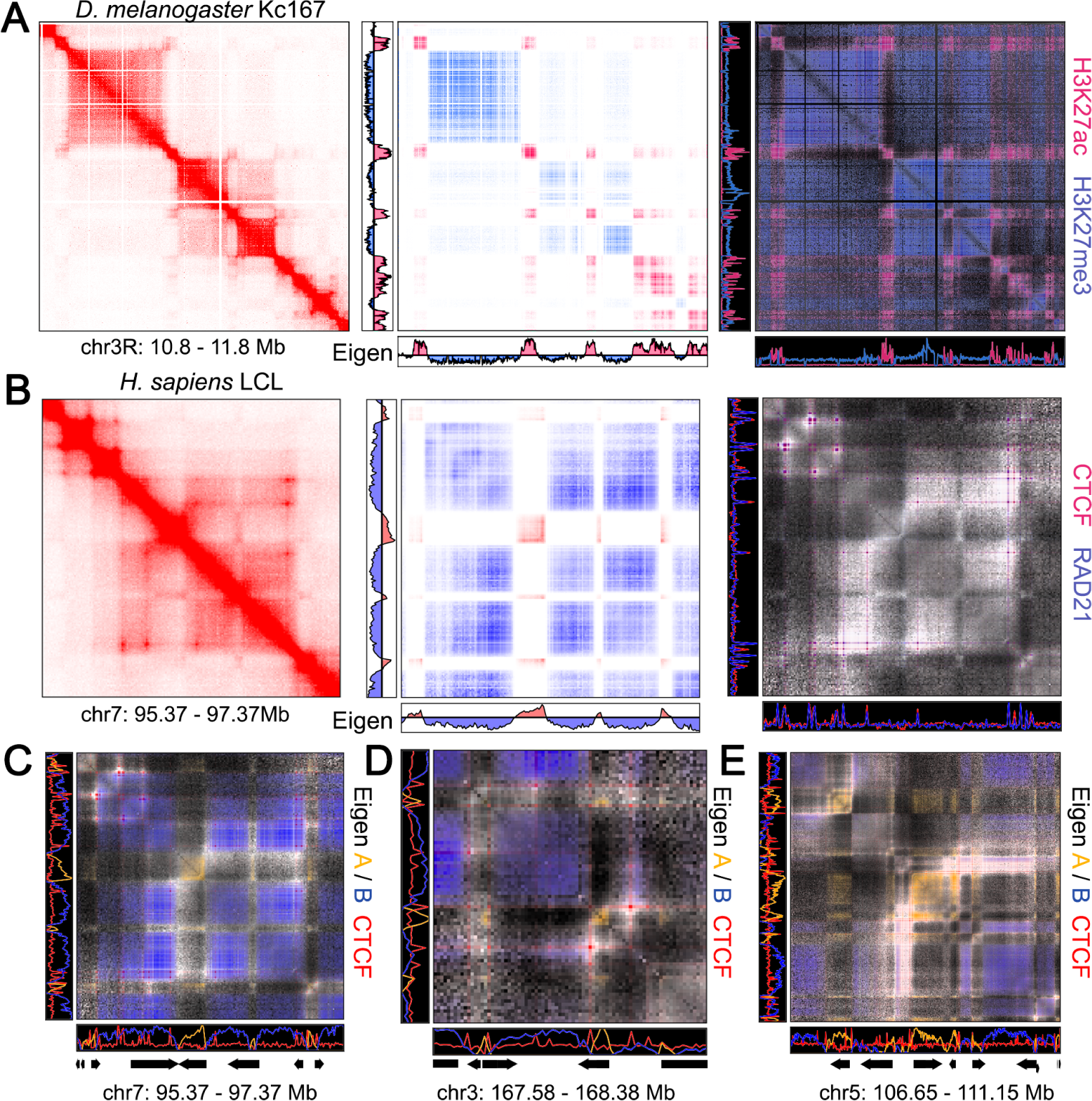
Interaction domains can be formed from different layers of chromatin organization. **A:** Domains of interactions in *D. melanogaster* Kc167 cells found by Hi-C (left), correspond to compartment intervals as colored by HiCrayon (middle), and alternating chromatin activity states (right) which is shown using HiCrayon coloring of H3K27ac (red) and H3K27me3 (blue). **B:** A 2 Mb region in human lymphoblastoid cell lines (LCL) showing loops and small compartment intervals by Hi-C (left), which become more distinguishable by HiCrayon coloring of the A/B compartment eigenvector (middle), and by HiCrayon coloring of ChIP-seq (right) for CTCF (red) and RAD21 (blue). **C-E:** HiCrayon coloring the A (orange) and B (blue) compartment eigenvector simultaneous with CTCF ChIP-seq (red).

We then examined near diagonal interactions in the recently published high-resolution Hi-C map of human lymphoblastoid cell lines (LCL), which showed that compartment intervals can be smaller than CTCF loops [5]. HiCrayon visualization of the 1 kb resolved compartment eigenvector, alongside CTCF and RAD21 ChIP-seq, effectively demonstrates that small compartment intervals can be smaller than CTCF loops and can segregate loci inside the loop (Figure 6B). Coloring A compartment (orange), B compartment (blue), and CTCF (red) interactions together in the same map helps to fully visualize the distinct organizational layers imparted by CTCF loops and compartments (Figure 6C). Examples of this at other loci reveal strongly interacting promoters in the A compartment, which interact despite crossing a CTCF loop boundary (Figure 6D). These promoter interactions can be particularly strong for genes proximal to each other in the A compartment (Figure 6E). Overall, this new visualization method reveals distinct layers that contribute to 3D genome organization.

## 3 Discussion

HiCrayon represents a novel way of co-visualizing 2D and 1D ‘omic datasets and is effective at detecting precise overlaps and distinguishing distinct features of genome organization. Using a simple weighting of 2D signals by 1D features, we demonstrate the utility of HiCrayon for visualizing different aspects of chromatin organization, such as punctate loops and compartments. While this approach can reveal previously difficult-to-visualize aspects of chromatin organization, such as multi-state compartments [41], there are limitations. For example, due to interpolation of 1D signal in 2D, it’s possible that HiCrayon could downplay an interaction that has ChIP-seq signal only at one anchor. However, these loci are likely rare or artificial events, especially considering that cross-linking can result in immunoprecipitation of interacting sites, a feature which is taken advantage of by methods like HiChIP and ChIA-PET [47–49].

We demonstrate several recently discovered features of chromatin organization, as well as some that are often debated. For example, we show a clear relationship between TADs, chromatin state, and compartments in *D. melanogaster* (Figure 6A), but other factors are likely involved in either forming or influencing these structures [4, 16, 18, 43, 45, 46]. Indeed, despite the overlap of TADs with compartment intervals in *D. melanogaster*, depletion of insulator proteins can result in altered genome organization [43, 45]. 3D chromatin organization is a complex interplay of multiple governing principles, and it is likely that TADs in *D. melanogaster* can represent features that exist from both the mechanisms that create compartment interactions and the activities of various architectural proteins.

As we demonstrate, HiCrayon visualization of the eigenvector helps distinguish A and B compartment interactions. However, it can also help identify interaction patterns that are more complex. This feature is useful both to visually evaluate the effectiveness of compartment identification [24] and to identify loci that don’t quite fit with the two-state A/B compartment model such as a 3rd compartment or, potentially, fairly distinct subcompartments [7, 41, 50–54]. Notably, when we color maps by marks of euchromatin, polycomb repressive chromatin, and heterochromatin, we obtain a more complete picture of the predominant interacting loci. However, we also note that not all of the interactions can be colored by just those three marks, suggestive of an even more complex compartmental organization.

## 4 Conclusions

We present HiCrayon, a tool to integrate 2D and 1D ‘omic data. We found that HiCrayon distinguishes 2D features that overlap distinct protein-bound loci, improving the precision at which these relationships can be visually explored. The use of HiCrayon to compare samples, such as maps of the differentiation process, allows a visual representation of the coordinated changes in protein occupancy with that of altered genome folding. HiCrayon’s ability to color by ChIP-seq and by compartment status allows greater exploration of a multiple-state model of compartmental organization. We found that it can also delineate near-diagonal interactions formed by distinct mechanisms, such as domains attributable to compartmental domains vs. CTCF loops. Furthermore, HiCrayon is an application designed with ease of use and reproducibility in mind with the ability to be used as a web or locally-hosted application to aid in production of publication-quality images.

## 5 Methods

### 5.1 Data

The lymphoblastoid Hi-C map used in Figures 2 and 6 is a combination of 8 public LCLs deposited in ENCODE or GEO repositories as follows: AK1: ENCSR508EMN, GM12878: ENCSR410MDC, GSE63525, GM12891: ENCSR859YSL, GM12892: ENCSR075VWI, GM13976: ENCSR634FNY, GM13977: ENCSR261EVH, GM18526: ENCSR693CIM, GM19239: ENCSR264SMC. Raw fastq files were downloaded and reprocessed to align to the human genome build hg38, using juicer v1.14.08 [19] with a quality filter of 30. The original fastq files, individual processed hg38 .hic files, and the combined 10 billion intrachromosomal contact Hi-C LCL map (.hic) are available at GEO accession GSE255264.

All Hi-C and ChIP-seq datasets used in this study are publicly available (Table 1). The eigenvector denoting compartments was identified in each of the corresponding Hi-C datasets using POSSUMM [5]. Hi-C distance normalization was performed as described previously [55] by taking the average signal at each diagonal as an expected value, in the following formula.

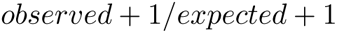

**Table 1.**
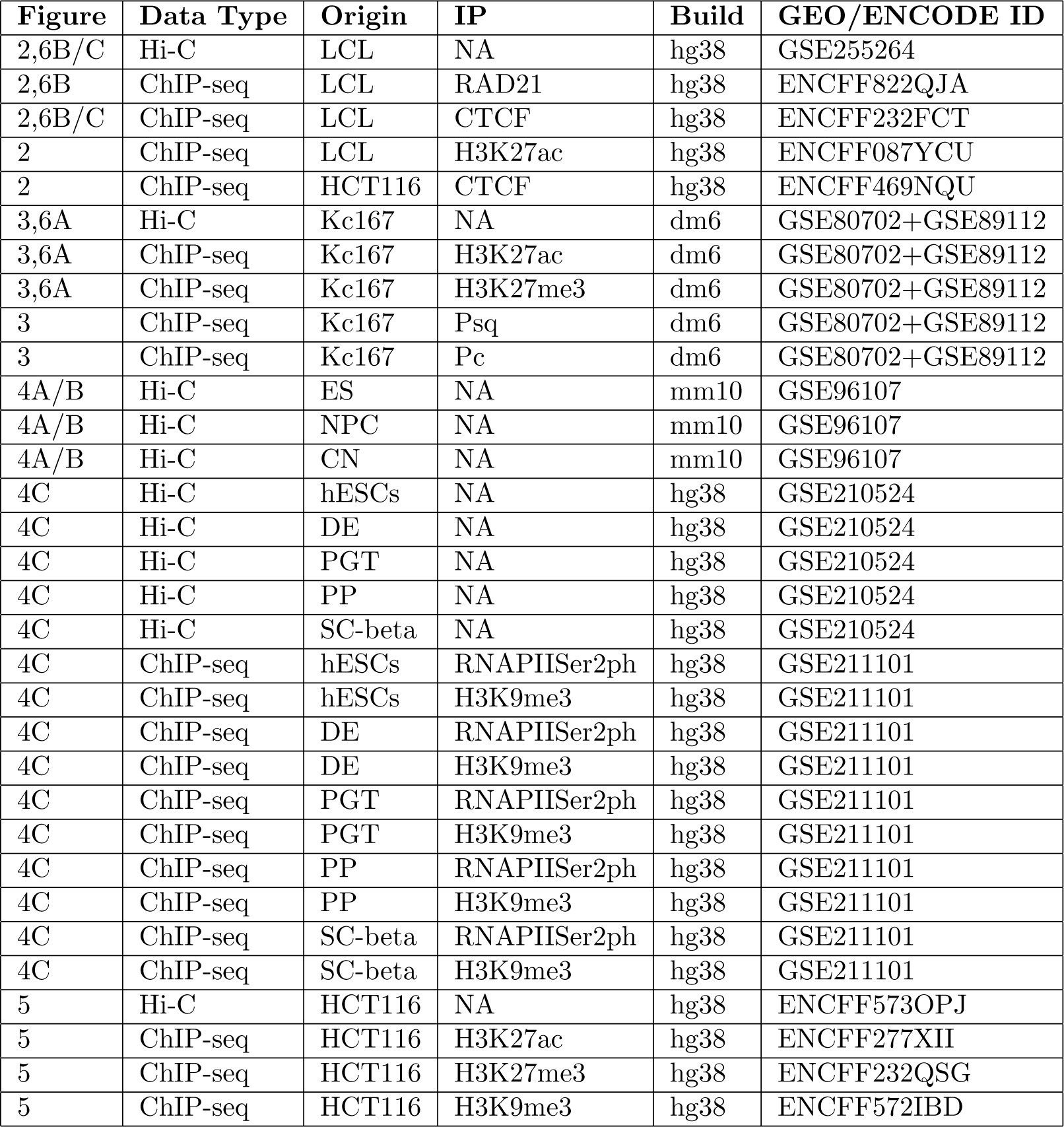
Accessions for publicly available data used in this paper.

### 5.2 Matrix Generation

HiCrayon generates a 2-dimensional matrix from a 1-D ChIP-seq track or a compartment call track. The resulting matrix is then weighted by contact frequencies derived from chromatin contact maps, such as a Hi-C contact matrix. HiCrayon streams contact frequencies from either a local file or URL using straw, storing the 2D contact matrix for a user-specified region. 1D signals in bigwig format (i.e. ChIP-seq) or bed-graph format (i.e. compartmental eigenvector) are then used to calculate a 1D to 2D signal matrix. First, 1D signal undergoes log transformation and scaling to fit all values between 0 and 1. In the case of 1D tracks that have negative values (i.e. the compartmental eigenvector), negative values are considered separately throughout the calculations and are multiplied by -1 before log transformation and scaling. Next, we create a two-dimensional signal matrix *m* where each value is the multiplicative product of the scaled 1D signal *s* in the row and column.

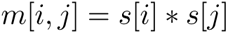

Separately, we also scale the 2D contact map between 0 and 1, after which the scaled contact matrix *c* is multiplied against the 1D signal matrix *m* and then converted to an 8-bit color scale to produce *h*:

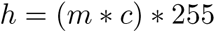

Finally, matrix *h* is adjusted to the user’s desired color scheme (RGB), resulting in an RGBA value for each bin within the matrix. These calculations result in an image where the transparency is a function of the joint ChIP-seq or compartment call data and contact intensity.

### 5.3 Color Blending

HiCrayon allows the overlay of multiple 1D tracks on the contact matrix in order to visualize potentially overlapping features. For each 1D matrix, *m* is generated as described above. We then combine these matrices using linear interpolation, a method that determines the intermediate value within the range of discrete values. In the context of color, linear interpolation finds an intermediate between *n* colors. Let *M_b_* be the *b*-th input matrix, where *b* = 1, 2*,…, n*, and *n* is the number of 1D signal matrices *m* in the input list. This allows the user to select *n* ChIP-seq tracks to visualize in a single matrix.

The alpha values for each matrix are then summed.

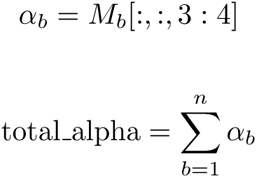

The blend ratios for each matrix are computed as

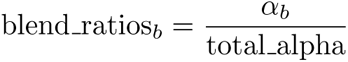

The color channels for each matrix are extracted as

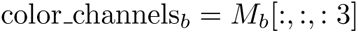

and the mixed-color channels are calculated as

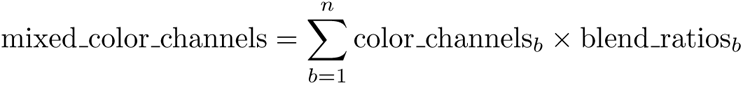

The final alpha value is limited to the 8-bit maximum:

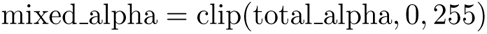

The final mixed matrix is then obtained by stacking the mixed color channels with the mixed alpha value.

## Acknowledgments

This work was supported by funding provided by the National Institutes of Health (NIH) Maximizing Investigators’ Research Award (MIRA) R35GM147467 to M.J.R. The content is solely the responsibility of the authors and does not necessarily represent the official views of the NIH. Funding for open access charge: National Institutes of Health.

## **6** Declarations

### **6.1** Ethics approval and consent to participate

Not applicable

### **6.2** Consent for publication

Not applicable

### **6.3** Availability of data and materials

A full version of HiCrayon can be downloaded and run locally from https://github.com/JRowleyLab/HiCrayon. A scaled-down web version of HiCrayon is available at https://jrowleylab.com/HiCrayon. Hi-C (.hic) files for the individual and combined LCL [5] maps reprocessed to genome build hg38 are available from GSE255264. ChIP-seq in GM12878 (LCL) was used for CTCF ENCFF749HDD, RAD21 ENCFF416LQD, H3K27ac The high-resolution Hi-C maps of *D. melanogaster* Kc167 cells are available from GSE80702 [8] and GSE89112 [10]. Hi-C files for mouse genome build mm10 ES, NPC, and CN [39] are available from GSE161259. HiC for human genome build hg38 pancreatic differentiation [38] are available from GSE210524. Hi-C in HCT116 [36] genome build hg19 is available from GSE104333. See Table 1).

### **6.6** Authors’ contributions

B.N. led the development of the underlying tool, R shiny implementation, and web-based version. H.L.H. and M.J.R. conceptualized the project. H.L.H., A.C., and T.E.R. tested the tool and suggested improvements. M.J.R. supervised the project, and B.N., C.T.C., and M.J.R. wrote and edited the manuscript.

### 6.7 Acknowledgements

We wish to acknowledge Dr. Mamta Shukla for helpful discussions over the development of this project.

## References

[1] Lieberman-Aiden, E., et al. Comprehensive mapping of long-range interactions reveals folding principles of the human genome. Science 326, 289–293 (2009). URL https://www.science.org/doi/10.1126/science.1181369. Publisher: American Association for the Advancement of Science.

[2] Hsieh, T.-H. S., et al. Mapping Nucleosome Resolution Chromosome Folding in Yeast by Micro-C. Cell 162, 108–119 (2015). URL https://www.cell.com/cell/abstract/S0092-8674(15)00638-8. Publisher: Elsevier.

[3] Deshpande, A. S., et al. Identifying synergistic high-order 3D chromatin conformations from genome-scale nanopore concatemer sequencing. Nature Biotechnology 40, 1488–1499 (2022). URL https://www.nature.com/articles/s41587-022-01289-z. Number: 10 Publisher: Nature Publishing Group.

[4] Rowley, M. J. & Corces, V. G. Organizational principles of 3D genome architecture. Nature Reviews. Genetics 19, 789–800 (2018).

[5] Harris, H. L. et al. Chromatin alternates between A and B compartments at kilo-base scale for subgenic organization. Nature Communications 2023 14:1 14, 1–17 (2023). URL https://www.nature.com/articles/s41467-023-38429-1. Publisher: Nature Publishing Group.

[6] Goel, V. Y., Huseyin, M. K. & Hansen, A. S. Region Capture Micro-C reveals coalescence of enhancers and promoters into nested microcompartments. Nature Genetics 55, 1048–1056 (2023).

[7] Rao, S. S. P. et al. A 3D map of the human genome at kilobase resolution reveals principles of chromatin looping. Cell 159, 1665–1680 (2014).

[8] Cubeñas-Potts, C., et al. Different enhancer classes in Drosophila bind distinct architectural proteins and mediate unique chromatin interactions and 3D architecture. Nucleic Acids Research 45, 1714–1730 (2017).

[9] Ogiyama, Y., Schuettengruber, B., Papadopoulos, G. L., Chang, J.-M. & Cavalli, G. Polycomb-Dependent Chromatin Looping Contributes to Gene Silencing during Drosophila Development. Molecular Cell 71, 73–88.e5 (2018).

[10] Eagen, K. P., Aiden, E. L. & Kornberg, R. D. Polycomb-mediated chromatin loops revealed by a subkilobase-resolution chromatin interaction map. Proceedings of the National Academy of Sciences of the United States of America 114, 8764–8769 (2017).

[11] Gutierrez-Perez, I. et al. Ecdysone-Induced 3D Chromatin Reorganization Involves Active Enhancers Bound by Pipsqueak and Polycomb. Cell Reports 28, 2715–2727.e5 (2019).

[12] Dixon, J. R. et al. Topological domains in mammalian genomes identified by analysis of chromatin interactions. Nature 485, 376–380 (2012).

[13] Sexton, T. et al. Three-dimensional folding and functional organization principles of the Drosophila genome. Cell 148, 458–472 (2012).

[14] Hou, C., Li, L., Qin, Z. S. & Corces, V. G. Gene density, transcription, and insulators contribute to the partition of the Drosophila genome into physical domains. Molecular Cell 48, 471–484 (2012).

[15] Nora, E. P. et al. Spatial partitioning of the regulatory landscape of the X-inactivation centre. Nature 485, 381–385 (2012).

[16] Rowley, M. J. et al. Evolutionarily Conserved Principles Predict 3D Chromatin Organization. Molecular Cell 67, 837–852.e7 (2017).

[17] Dong, P. et al. 3D Chromatin Architecture of Large Plant Genomes Determined by Local A/B Compartments. Molecular Plant 10, 1497–1509 (2017).

[18] Beagan, J. A. & Phillips-Cremins, J. E. On the existence and functionality of topologically associating domains. Nature Genetics 52, 8–16 (2020).

[19] Durand, N. C. et al. Juicebox Provides a Visualization System for Hi-C Contact Maps with Unlimited Zoom. Cell Systems 3, 99–101 (2016). URL 10.1016/j.cels.2015.07.012.

[20] Kerpedjiev, P. et al. HiGlass: web-based visual exploration and analysis of genome interaction maps. Genome Biology 19, 125 (2018).

[21] Chang, W., et al. shiny: Web application framework for R. manual (2023). URL https://shiny.posit.co/.

[22] ENCODE Project Consortium. An integrated encyclopedia of DNA elements in the human genome. Nature 489, 57–74 (2012).

[23] Luo, Y. et al. New developments on the Encyclopedia of DNA Elements (ENCODE) data portal. Nucleic Acids Research 48, D882–D889 (2020).

[24] Kalluchi, A., Harris, H. L., Reznicek, T. E. & Rowley, M. J. Considerations and caveats for analyzing chromatin compartments. Frontiers in Molecular Biosciences 10, 1168562 (2023). Publisher: Frontiers Media S.A.

[25] Hsieh, T.-H. S. et al. Resolving the 3D Landscape of Transcription-Linked Mammalian Chromatin Folding. Molecular Cell 78, 539–553.e8 (2020).

[26] Krietenstein, N. et al. Ultrastructural Details of Mammalian Chromosome Architecture. Molecular Cell 78, 554–565.e7 (2020).

[27] Nichols, M. H. & Corces, V. G. A CTCF Code for 3D Genome Architecture. Cell 162, 703–705 (2015).

[28] Alipour, E. & Marko, J. F. Self-organization of domain structures by DNA-loop-extruding enzymes. Nucleic Acids Research 40, 11202–11212 (2012).

[29] Nasmyth, K. Disseminating the genome: joining, resolving, and separating sister chromatids during mitosis and meiosis. Annual Review of Genetics 35, 673–745 (2001).

[30] Sanborn, A. L. et al. Chromatin extrusion explains key features of loop and domain formation in wild-type and engineered genomes. Proceedings of the National Academy of Sciences of the United States of America 112, E6456–6465 (2015).

[31] Fudenberg, G. et al. Formation of Chromosomal Domains by Loop Extrusion. Cell Reports 15, 2038–2049 (2016).

[32] Guo, Y. et al. CRISPR Inversion of CTCF Sites Alters Genome Topology and Enhancer/Promoter Function. Cell 162, 900–910 (2015).

[33] Haarhuis, J. H. I. et al. The Cohesin Release Factor WAPL Restricts Chromatin Loop Extension. Cell 169, 693–707.e14 (2017).

[34] Wutz, G. et al. Topologically associating domains and chromatin loops depend on cohesin and are regulated by CTCF, WAPL, and PDS5 proteins. The EMBO journal 36, 3573–3599 (2017).

[35] Nora, E. P. et al. Targeted Degradation of CTCF Decouples Local Insulation of Chromosome Domains from Genomic Compartmentalization. Cell 169, 930– 944.e22 (2017).

[36] Rao, S. S. P. et al. Cohesin Loss Eliminates All Loop Domains. Cell 171, 305–320.e24 (2017).

[37] Gong, H., Yang, Y., Zhang, S., Li, M. & Zhang, X. Application of Hi-C and other omics data analysis in human cancer and cell differentiation research. Computational and Structural Biotechnology Journal 19, 2070–2083 (2021).

[38] Lyu, X., Rowley, M. J., Kulik, M. J., Dalton, S. & Corces, V. G. Regulation of CTCF loop formation during pancreatic cell differentiation. Nature Communications 14, 6314 (2023).

[39] Bonev, B. et al. Multiscale 3D Genome Rewiring during Mouse Neural Development. Cell 171, 557–572.e24 (2017).

[40] Vilarrasa-Blasi, R. et al. Dynamics of genome architecture and chromatin function during human B cell differentiation and neoplastic transformation. Nature Communications 12, 651 (2021).

[41] Nichols, M. H., Correspondence, V. G. C. & Corces, V. G. Principles of 3D compartmentalization of the human genome. Cell Reports 35 (2021). URL 10.1016/j.celrep.2021.109330.

[42] Siegenfeld, A. P. et al. Polycomb-lamina antagonism partitions heterochromatin at the nuclear periphery 13, 4199. URL https://www.nature.com/articles/s41467-022-31857-5.

[43] Arzate-Mejía, R. G., Josué Cerecedo-Castillo, A., Guerrero, G., Furlan-Magaril, M. & Recillas-Targa, F. In situ dissection of domain boundaries affect genome topology and gene transcription in Drosophila. Nature Communications 11, 894 (2020).

[44] Ulianov, S. V. et al. Order and stochasticity in the folding of individual Drosophila genomes. Nature Communications 12, 41 (2021).

[45] Kaushal, A. et al. CTCF loss has limited effects on global genome architecture in Drosophila despite critical regulatory functions. Nature Communications 12, 1011 (2021).

[46] Ulianov, S. V. et al. Active chromatin and transcription play a key role in chromosome partitioning into topologically associating domains. Genome Research 26, 70–84 (2016).

[47] Skene, P. J. & Henikoff, S. An efficient targeted nuclease strategy for high-resolution mapping of DNA binding sites. eLife 6, e21856 (2017).

[48] Mumbach, M. R. et al. HiChIP: efficient and sensitive analysis of protein-directed genome architecture. Nature Methods 2016 13:11 13, 919–922 (2016). URL https://www.nature.com/articles/nmeth.3999. Publisher: Nature Publishing Group.

[49] Fullwood, M. J. & Ruan, Y. ChIP-based methods for the identification of long-range chromatin interactions. Journal of Cellular Biochemistry 107, 30–39 (2009).

[50] Yaffe, E. & Tanay, A. Probabilistic modeling of Hi-C contact maps eliminates systematic biases to characterize global chromosomal architecture. Nature Genetics 43, 1059–1065 (2011).

[51] Xiong, K. & Ma, J. Revealing Hi-C subcompartments by imputing interchromosomal chromatin interactions. Nature Communications 10, 5069 (2019).

[52] Wen, Z. et al. Extensive Chromatin Structure-Function Associations Revealed by Accurate 3D Compartmentalization Characterization. Frontiers in Cell and Developmental Biology 10, 845118 (2022).

[53] Ashoor, H. et al. Graph embedding and unsupervised learning predict genomic sub-compartments from HiC chromatin interaction data. Nature Communications 11, 1173 (2020).

[54] Liu, Y. et al. Systematic inference and comparison of multi-scale chromatin sub-compartments connects spatial organization to cell phenotypes. Nature Communications 12, 2439 (2021).

[55] Rowley, M. J. et al. Analysis of Hi-C data using SIP effectively identifies loops in organisms from C. elegans to mammals. Genome Research 30, 447–458 (2020).

